# Dynamic modelling of hepatitis C transmission among people who inject drugs: A tool to support WHO elimination targets

**DOI:** 10.1101/460550

**Authors:** Theresa Stocks, Leah J. Martin, Sharon Kühlmann-Berenzon, Tom Britton

## Abstract

To reach the WHO goal of hepatitis C elimination, it is essential to identify the number of people unaware of their hepatitis C virus (HCV) infection and to investigate the effect of interventions on the disease transmission dynamics. In many middle- and high-income countries, one of the primary routes of HCV transmission is via contaminated needles shared by people who inject drugs (PWIDs). However, substantial underreporting combined with high uncertainty regarding the size of this difficult to reach population, makes it challenging to estimate the core indicators recommended by the WHO. To help enable countries to monitor their progress towards the elimination goal, we present a novel multi-layered dynamic transmission model for HCV transmission within a PWID population. The model explicitly accounts for disease stage (acute and chronic), injection drug use status (active and former PWIDs), status of diagnosis (diagnosed and undiagnosed) and country of disease acquisition (domestic or abroad). First, based on this model, and using routine surveillance data, we estimate the number of undiagnosed PWIDs, the true incidence, the average time until diagnosis, the reproduction numbers and associated uncertainties. Second, we examine the impact of two interventions on disease dynamics: 1) direct-acting antiviral drug treatment, and 2) needle exchange programs. To make the model accessible to relevant users and to support communication of our results to public health decision makers, the model and its output are made available through a Shiny app. As a proof of concept, we illustrate our results for a specific data set; however, through the app our model can be easily adapted to other high-income countries with similar transmission patterns among PWIDs where the disease is endemic.

## 1 Introduction

Hepatitis C virus (HCV) is a bloodborne pathogen which can, left untreated, lead to chronic hepatitis, cirrhosis, liver cancer and death (Heymann, 2004). Worldwide, an estimated 71 million people are chronically HCV-infected, causing approximately 399 000 deaths each year; however, only an estimated 20% are aware of their infections (WHO, 2018a). In many middle- and high-income countries, injection drug use is the leading cause of HCV transmission, via sharing of contaminated needles (Hajarizadeh et al., 2013). As no vaccine against HCV is currently available, treatment as well as harm-reduction approaches like needle exchange programs (NEPs) for people who inject drugs (PWIDs) are critical to reducing prevalence and incidence. With recently introduced direct acting antivirals (DAAs), up to 95% of those treated achieve HCV treatment success and are no longer infected or infectious (Lawitz et al., 2014; Dore and Jordan, 2015). The World Health Organization (WHO) recommends treatment with DAA-based therapies for all HCV-infected individuals, with some specific exceptions based on genotype (WHO, 2018a). However, access to diagnosis and treatment remains limited; in 2015, only 7% of all diagnosed cases worldwide received treatment (WHO, 2018a). NEPs offer PWIDs the opportunity to replace used syringes, needles and injection tools with new sterile ones in order to reduce transmission of HIV, hepatitis B and C, and other bloodborne pathogens. Moreover, these programs provide HIV and hepatitis testing and linkage to treatment and are also associated with education, counseling and referral to substance use disorder treatment (CDC, 2018).

Following the introduction of DAAs, the member states of the World Health Assembly, which governs the WHO, have committed to reducing viral hepatitis mortality by 65% and incidence by 90% (using 2015 as a baseline) by 2030 (WHO, 2018b). Two of the core indicators that countries are advised to monitor to measure progress towards this goal are: a) HCV prevalence (indicator C.1.b in (WHO, 2018b)), including both diagnosed and undiagnosed cases, and b) HCV incidence (indicator c.9.b in (WHO, 2018b)); both measures should be disaggregated by risk group, including PWIDs. Prevalence surveys are a preferable source of data for these estimates but such studies are usually costly to perform and the population of interest, in particular PWIDs, is often not easy to reach. Mathematical models can be used to complement these approaches (WHO, 2018b) and are useful tools that can generate scientific evidence in order to guide policies (Scott et al., 2018b).

The primary objective of this study is to develop a dynamic transmission model to help enable country-specific estimation of four key quantities and associated uncertainties to monitor progress towards the WHO HCV elimination goals among PWIDs: 1) the number of undiagnosed cases, 2) true incidence of HCV, i.e. the annual number newly infected (as opposed to newly diagnosed) individuals, 3) average time until diagnosis and 4) reproduction numbers from routinely collected surveillance data. The second objective of this work is to examine the effects that DAA treatment and NEPs may have on prevalence among active PWIDs, true incidence, number of undiagnosed cases and total number of infected in the PWIDs risk group.

Modelling of HCV transmission among PWIDs and the impact of interventions via compartmental models has been studied before, e.g. Martin et al. (2011, 2013); Gountas et al. (2017) and Fraser et al. (2018). Often, HCV transmission models implicitly assume that everyone in the PWID population is currently injecting drugs (active) and that everyone who is infected is diagnosed. This can produce misleading results for two reasons. First, routine HCV surveillance data usually does not distinguish between former and active PWIDs. Therefore, calibrating such models to routine surveillance data can create the illusion of a larger active, and transmitting, PWID population than actually exists because former PWIDs are counted as infectious as well. Second, by ignoring undiagnosed cases, such models assume a smaller HCV-infected population than actually exists. By extension, assuming that all cases are diagnosed will lead to the mistaken assumption that all cases will have access to treatment. Recently, HCV transmission models stratified by HCV-diagnosis and injection drug use status have been proposed in order to investigate the effect of testing and treatment as prevention, e.g. Scott et al. (2017a,b, 2018a,b).

We demonstrate our model and the respective results calibrated to surveillance data from Sweden and other literature-informed values, however our approach goes beyond analyzing a specific data set. Rather we develop a general web-based tool (Shiny app), available for other countries with a similar disease situation represented in this work, to calibrate the model to their available data and, in that way, monitor their country-specific situation *and* investigate the effect of different intervention strategies. It is important to make our model accessible to relevant users, including public health decision-makers, to bridge the gap between theoretical models and real public health applications.

## 2 Methods

We develop a dynamic, deterministic ODE-based compartmental model to describe HCV transmission among PWIDs stratified by stage of infection (acute or chronic), injection drug use status (active or former PWIDs), status of diagnosis (undiagnosed or diagnosed), country of disease acquisition (within or outside the country under consideration). We consider two interventions: participation in NEPs (yes or no) and treatment with DAA (yes or no). We limit our study to the PWID population and we assume that the disease is only transmitted by sharing of contaminated needles within this community; HCV cases with other routes of transmission are not considered here.

We begin by describing the general underlying transmission dynamics in the base model and then describe how the two interventions are incorporated into the model. By definition, we assume that only active PWIDs can enroll in NEPs and that only diagnosed PWIDs can be treated. The schematic representation of the model is given in the flow diagram (Figure 1) where arrows indicate the movement between the compartments, with a list of notations (Table 1). The transmission equations and further mathematical details of the model are given in Section A of the Appendix.

**Figure 1:**
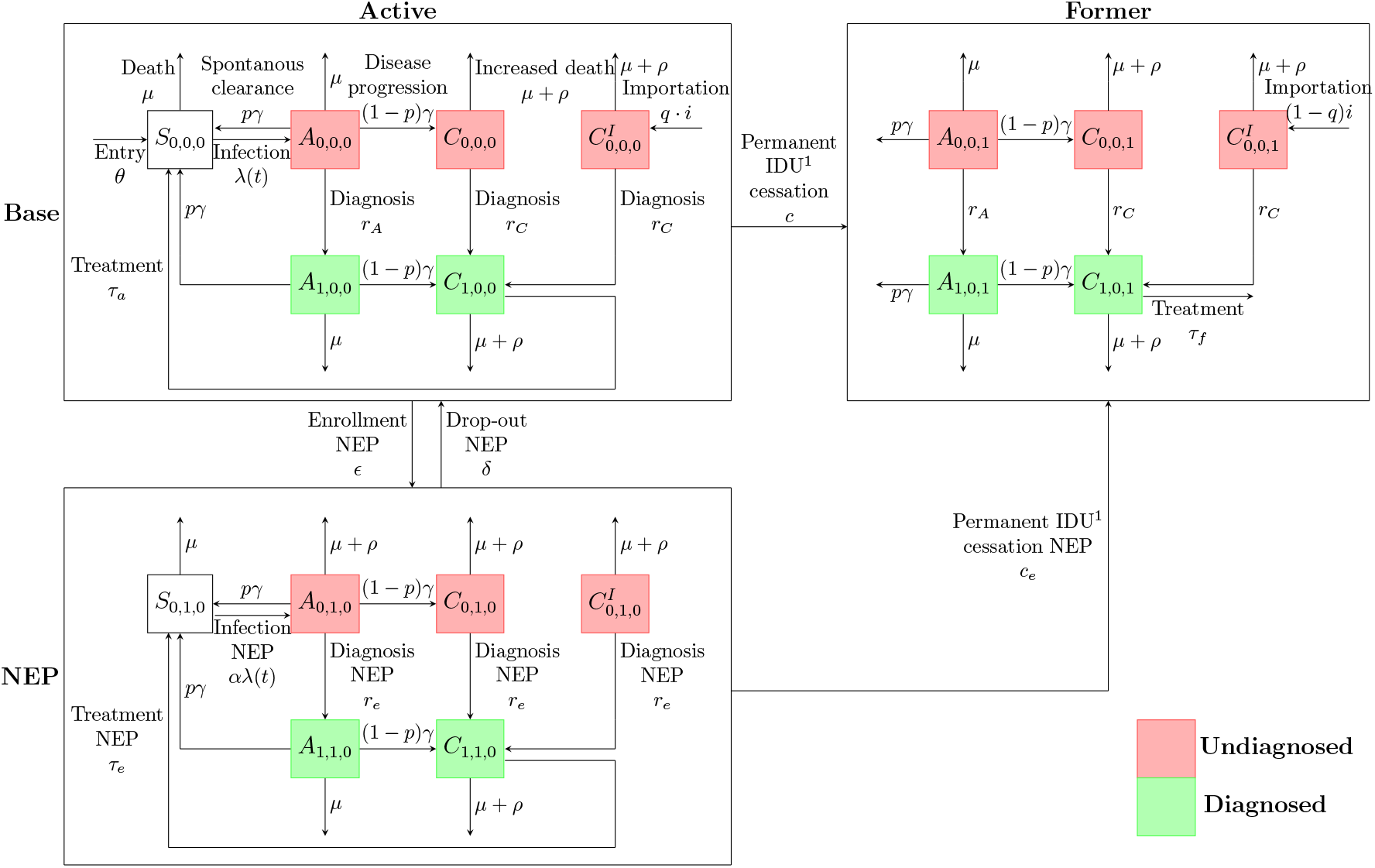
Schematic representation of the base transmission model and the needle exchange program (NEP). We distinguish between susceptible (*S_d,e,f_*) and two stages of infection, acute (*A_d,e,f_*) and chronic (*C_d,e,f_*). The subindices indicate *d* = diagnosed, *e* = enrolled in NEP and *f* = former PWIDs, where 0=no and 1=yes, and super-scripted with *I* if imported. The rates on the arrows are explained in the text. Injection drug use

**Figure 2:**
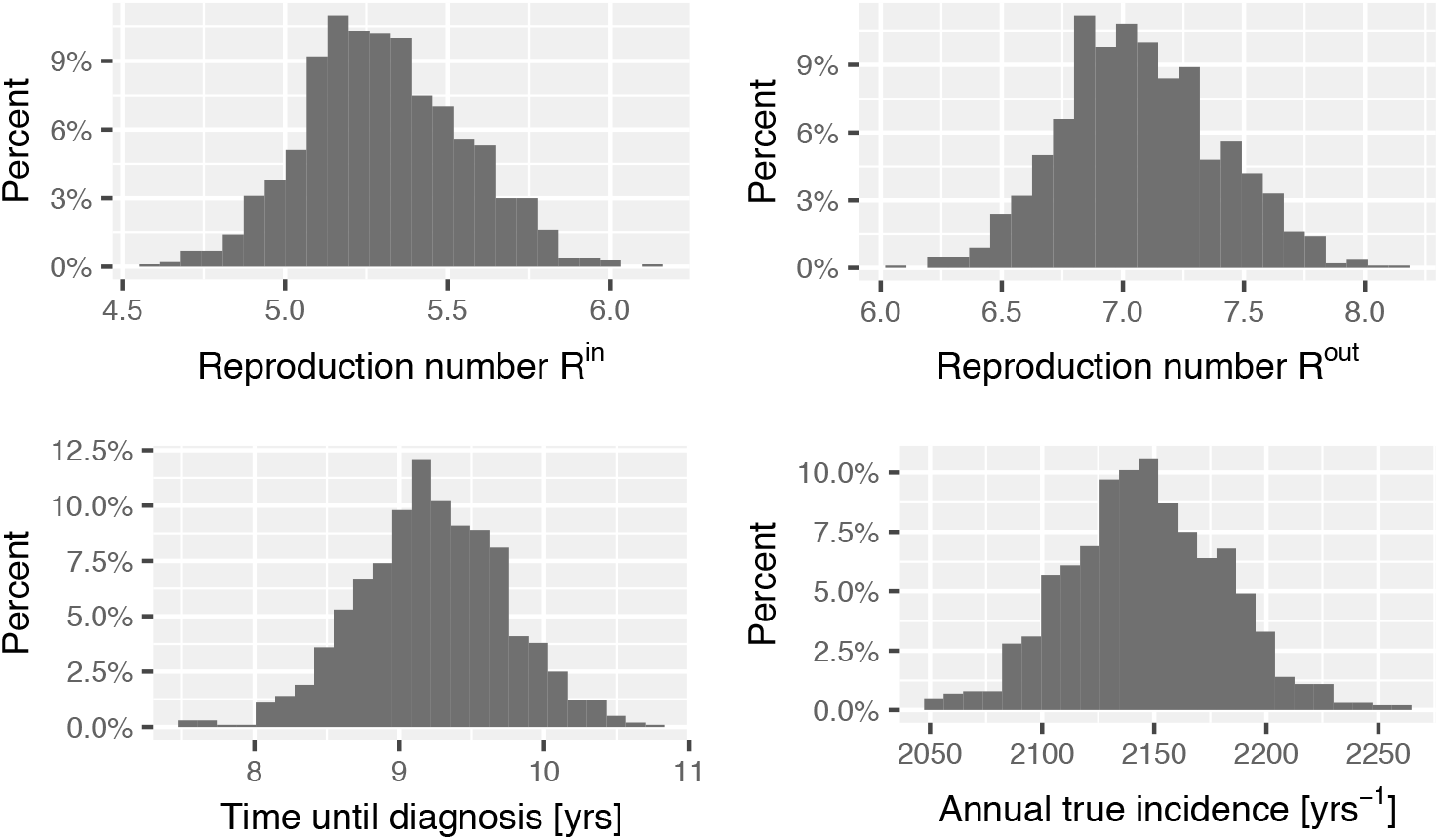
Distribution of the reproductions numbers *R*^in^ and *R*^out^, the time until diagnosis and the annual true incidence.

### 2.1 Base transmission model without interventions

The underlying basic transmission dynamics assume an SI-epidemic model (Keeling and Rohani, 2008) where susceptible individuals, once infected, move to an acute phase and eventually progress to become chronically-infected individuals, a state from which recovery is not possible without treatment. We assume that the active PWID population is constant over time. PWIDs can enter this population as either susceptible or imported (i.e. not domestically-acquired) chronically HCV-infected and leave by either death or permanent injection drug use cessation, the latter thus reclassifies them as former PWIDs. Active PWIDs can be infected only by other active infected PWIDs via sharing of contaminated needles while former PWIDs do not contribute to the infectious pressure. Once infected, PWIDs are initially undiagosed and may eventually progress to become diagnosed, which is when they are observed in the surveillance system as part of the annual reported incidence.

#### Compartments

We denote by *S_d,e,f_*(*t*) the number of susceptible PWIDs at time *t*, by *A_d,e,f_*(*t*) and *C_d,e,f_*(*t*) the number of acute and chronically infected PWIDs at time *t* and by 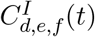 the number of imported, chronically infected PWIDs at time *t*. The indices denote diagnosis status (undiagnosed or **d**iagnosed, *d* = 0 or 1), enrollment in a NEP (no exchange or **e**xchange, *e* = 0 or 1) and injection drug use status (active or **f**ormer, *f* = 0 or 1). Since the model including interventions builds upon the base model, we include an index to indicate enrollment in the NEP, however, this index in the base model is set to *e* = 0 in all compartments. Note that some combinations of indices are not defined because they are not possible, e.g. *S*_1,0,0_(*t*), meaning the number of diagnosed susceptible individuals, because diagnosis requires infection. Altogether our model consists of 17 compartments out of 4∙2^3^ = 32 possible index combinations.

#### Entering the (active) PWID population

New PWIDs enter the active PWID population by two routes: either initially susceptible to HCV infection, at a rate *θ*, or as imported, undiagnosed, chronically-infected PWIDs, at a rate *q · i*. The influx rate *θ* into the susceptible compartment *S*_0,0,0_(*t*) is adjusted so that, in a system without interventions, the population size of active PWIDs, *N*, is constant over time, see Equation (13) in the Appendix.

#### Force of infection and disease progression

Susceptible PWIDs become infected at a per capita rate *λ*(*t*). This force of infection is proportional to the background prevalence of infected active PWIDs, see Equation (1). In the acute phase a fraction, *p*, of individuals naturally recovers from the disease and returns to the susceptible compartment, while the remaining (1 *− p*) infected individuals proceed to the chronic phase (Micallef et al., 2006). The rate of becoming chronically infected from the acute phase is *pγ* and the rate of natural recovery is (1 *− p*)*γ*.

#### Exiting the PWID population

Active PWIDs leave the PWID population by either death or permanent injection drug use cessation. Active PWIDs stop injecting drugs and move to the former PWID compartments at an annual cessation rate, *c*. We assume that all susceptible and acutely infected individuals have an all-cause, injection drug use-specific, mortality rate *μ* regardless of the status of their injection drug use or whether or not they have been diagnosed. However, the model allows for an increased death rate, *μ* + *ρ* with *ρ* ≥ 0, due to the progression of the disease, for both active and former chronically-infected PWIDs (Hajarizadeh et al., 2013).

#### Imported cases

We assume that there is a constant importation of undiagnosed, chronically-infected PWIDs, accounting for individuals who acquire the disease through injection drug use from outside the country under consideration. We assume that importation of acute cases is negligible because the duration of the acute phase is short (Mondelli et al., 2005). Out of all imported cases *i*, a fraction 0 ≤ *q* ≤ 1, enters as active and another (1 *− q*) as former PWIDs. The fraction *q* is calibrated to ensure that the ratio of active to former PWIDs amongst imported cases is proportional to the number of years an individual remains in the active or former PWID compartments, respectively (Equation (14) in the Appendix). Other rates for imported PWIDs are assumed to be identical to those for PWIDs infected within the country under consideration.

#### Diagnosis rates

Undiagnosed infected individuals are diagnosed based on a rate dependent upon the stage of infection, acute (*r_A_*) or chronic (*r_C_*). However, we assume that diagnosis rates do not vary directly due to injection drug use status (active or former), i.e. active and former PWIDs are diagnosed at the same rate and is the same for imported cases and domestic cases.

### 2.2 Modelling of interventions

#### DAA treatment

We assume that only diagnosed, chronically-infected PWIDs are eligible for DAA treatment and that no treatment is provided during the acute stage of infection because this stage is comparably short and natural recovery is possible (WHO, 2018a). For simplicity we assume that once treatment is initialized the patient immediately recovers. This is a sensible assumption because successful DAA treatment is highly effective and treatment durations are relatively short (Lawitz et al., 2014). Treatment rates depend upon injection drug use status (active or former) and enrollment in a NEP. The parameters *τ_a_*, *τ_e_* and *τ_f_* denote successful treatment rates for active PWIDs not enrolled in NEPs, active PWIDs enrolled in NEPs and former PWIDs, respectively. Successfully treated active PWIDs return to the respective susceptible compartment while successfully treated former PWIDs leave the PWID population and, therefore, the model.

#### Needle exchange programs

We assume that active PWIDs enroll uniformly into NEPs at a rate *ε*, regardless of diagnostic status or stage of infection, and drop-out of the programs at a rate *δ*, returning to the respective compartments in the base model. The dynamics in a NEP can potentially differ from the base model in three ways and our model allows these to vary independently. First, NEPs provide access to clean needles and injection equipment; therefore, we assume that the risk of both transmitting HCV and being infected reduces by a factor 0 ≤ *α* ≤ 1 (*α* = 1 meaning no reduction) compared to the susceptibility and infectiousness of infected, active PWIDs in the base model. Second, the model provides the potential for an increased diagnosis rate, *r_e_*, because regular health check-ups that include HCV screening often occur in these programs, independenlty of stage of infection. As a consequence, *r_e_* might differ from the diagnosis rates in the base model. Third, due to support and education available through medical facilities associated with NEPs, the model allows for inclusion of an increased permanent injection drug use-cessation rate, *c_e_*, compared to the cessation rate *c* in the base model.

### 2.3 Force of infection

The force of infection is defined as

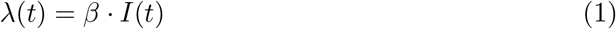

With

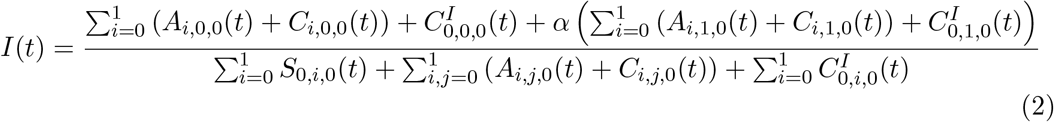

The denominator of *I*(*t*) is the sum of all active PWIDs while the numerator is the sum of all active and infected PWIDs with *α* accounting for the reduced infectiousness of PWIDs in NEPs. For *α* = 1, *I*(*t*) denotes the prevalence of HCV among the active PWID community at time *t*.

### 2.4 Derivation of key outcomes

#### 2.4.1 Number of undiagnosed PWIDs

From this model formulation we can now compute the number of diagnosed as well as undiagnosed at any time *t*, stratified by injection drug use status (active or former) and stage of infection (acute or chronic) which is the sum of the respective compartments at that time. The number of, e.g., undiagnosed, active, infected PWIDs at time *t* is equal to

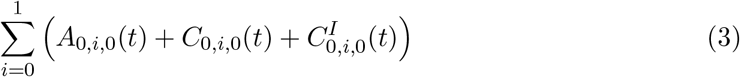

and the number of undiagnosed, former, infected PWIDs at time *t* is equal to

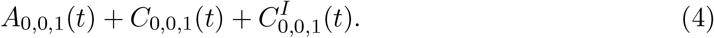

The total number of undiagnosed HCV-infected PWIDs in our model is then the sum of (3) and (4). The total number of infected (diagnosed and undiagnosed) PWIDs is the sum of all compartments except the two susceptible compartments.

#### 2.4.2 True incidence

The true incidence in year *t_n_* is the number of newly *infected* individuals in (*t_n−_*_1_*, t_n_*]. From the model the true annual incidence is given as

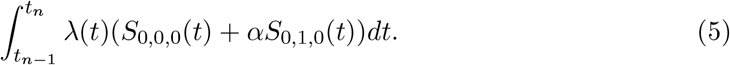

Note, that this is different from the reported incidence rate, which is the number of newly *diagnosed* individuals in year *t_n_*

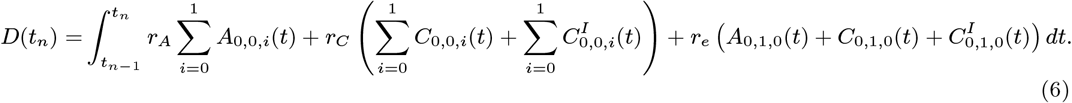

#### 2.4.3 Time until diagnosis

From the base model we can derive the average time from infection until diagnosis for a intervention-free system in endemic state. In the following, the endemic level is indicated by lowercase letters, e.g. in endemicity *A*_0,0,0_(*t*) = *a*_0,0,0_ for all *t*. Note that there is no disease-free state because we allow for importation of infectious individuals. In the system without interventions, the treatment rates, *τ_f_, τ_a_, τ_e_*, and the enrollment rate into NEPs, *ε*, are all set to zero. This represents a scenario in which both improved treatment options and broader access to NEPs are not available, as is the situation in many middle and high-income countries. The average time until diagnosis, *t*_Σ_, is given as the sum of the average time until diagnosis, conditional on the compartment within which an individual is diagnosed, weighted by the fraction of diagnosed in that respective compartment. Let *t_X_* denote the average time until diagnosis given the individual is diagnosed in compartment *X_d,e,f_* with *X* ∈ {*S, A, C*} (see Equations (15) of Section B in the Appendix). We then obtain

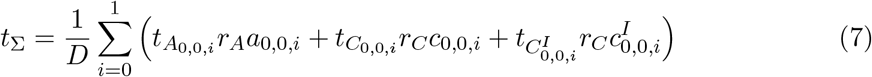

with *D* as in Equation (6) being the total number of newly diagnosed HCV cases each year for *t_n_ − t_n−_*_1_= 1 year in endemic, intervention-free state. Each infected individual will belong to the undiagnosed cohort from the time of infection up until the time of diagnosis. As a consequence, the number of undiagnosed in steady state will equal the incidence rate multiplied by the average time to diagnosis in steady state.

#### 2.4.4 Reproduction numbers *R*^in^ and *R*^out^

We define the reproduction numbers *R*^in^ and *R*^out^ as the expected number of new infections caused by a typical infectee who acquires the disease in the target country (in) or abroad (out), during the early stage of an epidemic when almost everyone is susceptible. They are calculated as the product of the number of individuals an infected individual infects on average per unit time and the duration of the infectious period. Since we assume that all infected individuals are equally infectious regardless of their stage of infection, we obtain the following for the base transmission model without interventions

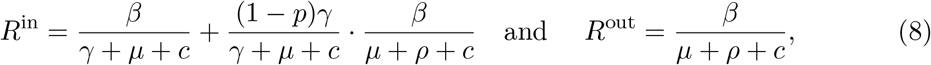

where the first factor in the second term in *R*^in^ is the probability of reaching the chronic state. Note that the reproduction numbers are not threshold parameters as long as there is importation of infectious individuals. However, without importation of infected PWIDs (i.e. *i* = 0), *R*^in^ = *R*_0_, where *R*_0_ denotes the basic reproduction number (Diekmann et al., 2013, Chapter 1, Page 4). Hence, if *R*^in^ ≤ 1 without importation, then no outbreak is possible. We illustrate this in Figure 3 in Section C in the Appendix.

**Figure 3:**
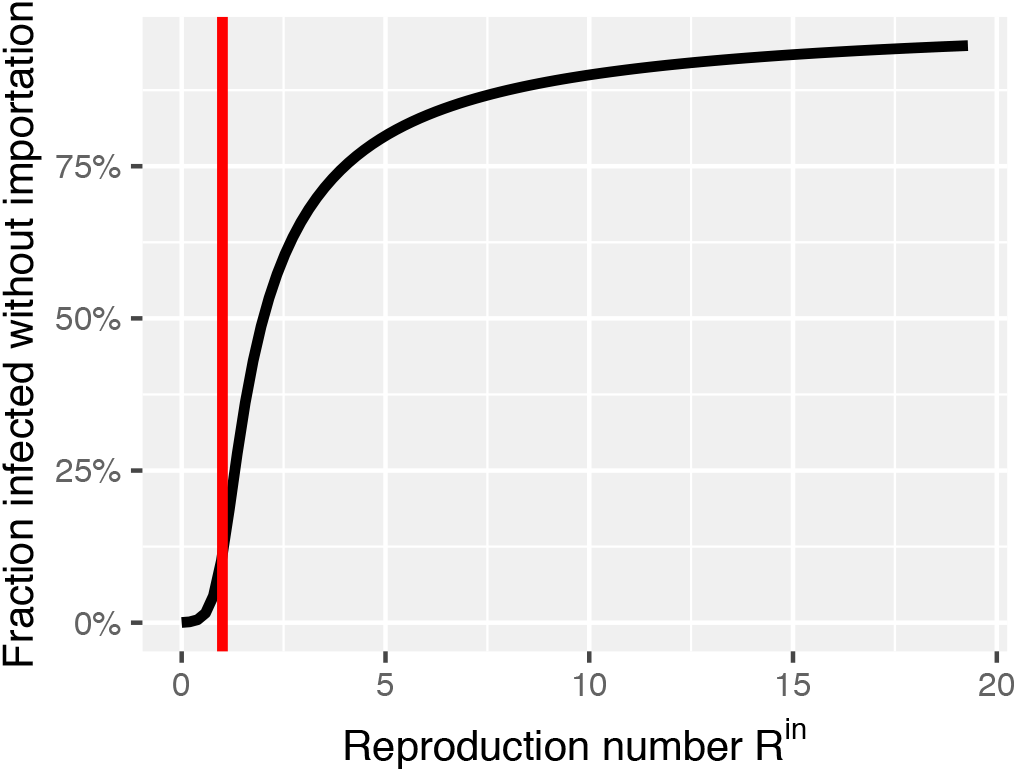
Fraction of infected active PWIDs dependent on *R*^in^ without importation.

**Figure 4:**
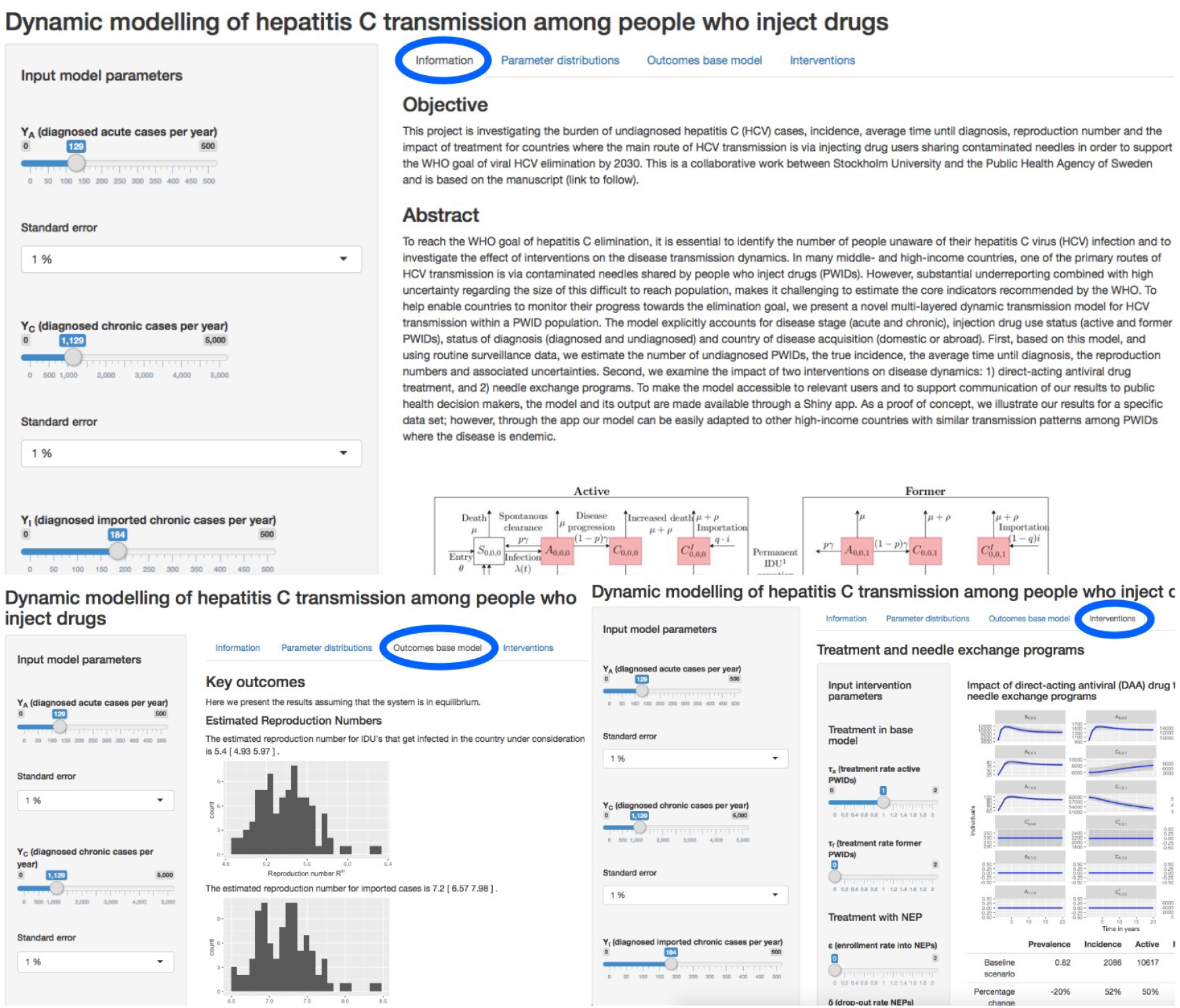
Screenshot of the shiny app.

### 2.5 Fitting the model to data

We assume HCV to be endemic before the implementation of interventions. For a given set of parameters, the endemic level is obtained by setting all derivatives in the system of equations in Section A in the Appendix to zero and solving the equations numerically; for this we use the R-package nleqslv (Hasselman, 2017). Our base model without interventions contains ten parameters of which we assume six to be known from the literature (Table 1). Moreover, we assume that the following four quantities are known: a) prevalence of HCV among active PWIDs in an intervention-free scenario, and the reported incidence of b) acute, c) chronic and d) imported, chronic cases. These four quantities can be formulated in terms of the model parameters (as explained below) and hence allow us to estimate the remaining four parameters, namely the reporting rates of acute and chronically infected PWIDs, *r_A_* and *r_C_*, the number of infectious contacts per time unit, *β*, and the importation rate *i*. Knowing all ten parameters in our model, we can then also calculate functions of the parameters such as the force of infection *λ*(*t*), the influx rate into the active PWID population *θ*, and the fraction of active chronically-infected imported cases *q*. We conduct a sensitivity analysis (Section 3.2) for all parameters that are assumed to be fixed and known.

**Table 1:** Notation and parameter values used in the model.

| Parameter | Explanation | Value | (sd) | [Unit] | Assumption | Reference |
| --- | --- | --- | --- | --- | --- | --- |
| **Compartments** |  |  |  |  |  |  |
| | $d$= diagnosed no/yes | {0,1} | | | | |
| | $e$= needle exchange program no/yes | {0,1} | | | | |
| | $f$= former PWIDs no/yes | {0,1} | | | | |
| $S_{d,e,f}(t)$ | Number of susceptible at time $t$ | | | [individuals] | | |
| $A_{d,e,f}(t)$ | Number of acutely infected at time $t$ | | | [individuals] | | |
| $C_{d,e,f}(t)$ | Number of chronically infected at time $t$ | | | [individuals] | | |
| $C_{d,e,f}^I(t)$ | Number of imported chronically infected at time $t$ | [individuals] | | | | |
| **Model without interventions** |  |  |  |  |  |  |
| $c$ | permanent injection drug use cessation rate | 0.056 | (0.00056) | [yrs<sup>-1</sup>] | known | (Kåberg et al., 2017) |
| $\mu$ | mortality rate for PWIDs | 0.022 | (0.00022) | [yrs<sup>-1</sup>] | known | (Mathers et al., 2013) |
| $\rho$ | factor by which mortality rate increases due to chronic HCV | 0 | (0) | | known | *not included in the model* |
| $p$ | spontaneous clearance probability | 0.26 | (0.0026) | | known | (Micallef et al., 2006) |
| $\gamma$ | rate of leaving acute compartment | 2 | (0) | [yrs<sup>-1</sup>] | known | (Mondelli et al., 2005) |
| $N$ | active PWID population size in endemic state | 26000 | (260) | [individuals] | known | (SWE, 2011, 2013, 2014) |
| $\beta$ | number of infectious contacts per time unit | 0.55 | (0.03) | [yrs<sup>-1</sup>] | calibrated | |
| $r_A$ | diagnosis rate of acutely infected PWIDs | 0.129 | (0.003) | [yrs<sup>-1</sup>] | calibrated | |
| $r_C$ | diagnosis rate of chronically infected PWIDs | 0.072 | (0.006) | [yrs<sup>-1</sup>] | calibrated | |
| $i$ | importation rate of active chronically infected cases | 53.02 | (1.2) | [individuals/yrs] | calibrated | |
| prev | HCV prevalence among active PWIDs before interventions | 0.817 | (0.00817) |  | Equation (10), known | (Fraser et al., 2018) |
| $Y_A$ | Diagnosed acute cases per year | 129 | (1.29) | [individuals/yrs] | Equation (9), known | Public Health Agency of Sweden |
| $Y_C$ | Diagnosed chronic cases per year | 1129 | (11.29) | [individuals/yrs] | Equation (9), known | Public Health Agency of Sweden |
| $Y_{C^I}$ | Diagnosed imported chronic cases per year | 184 | (1.84) | [individuals/yrs] | Equation (9), known | Public Health Agency of Sweden |
| $\lambda(t)$ | force of infection | | | [yrs<sup>-1</sup>] | Equation (1) | |
| $\theta$ | PWIDs recruitment rate | 1982 | (25) | [individuals/yrs] | Equation (13) | |
| $q$ | fraction of active imported chronic cases | 0.22 | (0) | | Equation (14) | |
| $R^{\text{in}}$ | reproduction number (infection within country) | 5.3 | (0.2) | | Equation (8) | |
| $R^{\text{out}}$ | reproduction number (infection outside country) | 7.1 | (0.3) | | Equation (8) | |
| $t_\Sigma$ | average time until diagnosis | 9.2 | (0.5) | [yrs] | Equation (7) | |
| **Treatment** |  |  |  |  |  |  |
| $\tau_a$ | treatment rate of active PWIDs (not in NEPs) | | | [yrs<sup>-1</sup>] | varied in Section 3.3 | |
| $\tau_e$ | treatment rate of active PWIDs in NEPs | | | [yrs<sup>-1</sup>] | varied in Section 3.3 | |
| $\tau_f$ | treatment rate of former PWIDs (not in NEPs) | | | [yrs<sup>-1</sup>] | varied in Section 3.3 | |
| **Needle exchange programs (NEPs)** |  |  |  |  |  |  |
| $r_e$ | diagnosis rate in NEPs | | | [yrs<sup>-1</sup>] | varied in Section 3.3 | |
| $\alpha$ | relative risk of acquiring/spreading HCV | | | | varied in Section 3.3 | |
| $c_e$ | permanent cessation rate of PWIDs in NEPs | | | [yrs<sup>-1</sup>] | varied in Section 3.3 | |
| $\epsilon$ | enrollment rate into NEPs | | | [yrs<sup>-1</sup>] | varied in Section 3.3 | |
| $\delta$ | drop-out rate NEPs | | | [yrs<sup>-1</sup>] | varied in Section 3.3 | |
| $m$ | sample size | 1000 | | | | |

**Table 2:**
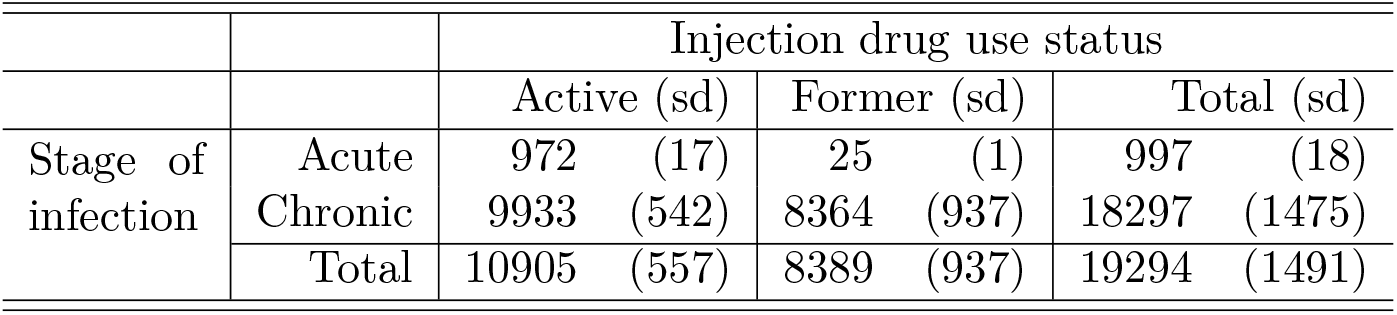
Mean number of estimated undiagnosed PWIDs cases (sd) stratified by stage of infection and status of drug use in steady state.

Our observations *Y_X,n_* are the number of newly diagnosed acute (A), chronic (C) and imported chronic (C^I^) HCV cases aggregated over reporting years (*t_n−_*_1_*, t_n_*], *n* ∈{1*, …, N*} of type *X* ∈ *{A, C, C^I^}*. Note that since reported surveillance data usually does not differentiate diagnosed individuals by injection drug use status (active or former), we only observe the sum of these two groups. Hence, for *t_n_ − t_n−_*_1_ = 1 year, the number of newly diagnosed during an interval of one year is given as

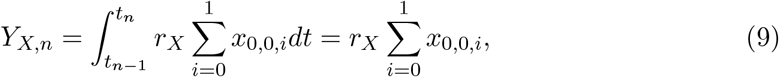

with *x* ∈ {*a, c, c^I^*} and 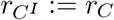. Since the observations are more or less the same each year when at endemic equilibrium, we only write *Y_X_* := *Y_X,n_* in what follows. HCV prevalence among active PWIDs in endemic state before interventions is given as

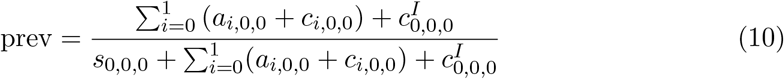

#### 2.5.1 Incorporating uncertainty

To account for uncertainty in the given parameters of the model, we assume that each literature-informed value is the mean of a (truncated) normal distribution bounded by zero from below and we assign an additional standard deviation (1% of the mean value in the numerical example) to these values (Table 1). We draw *m* samples of parameter values from these distributions assuming no dependence between parameter uncertainties. For each sampled parameter vector, we then solve the system of equations numerically and calculate the key outcomes of interest (Equations (3), (4), (5), (7), (8)). The reported estimated key outcomes are the resulting sample mean and the corresponding sample standard deviation.

#### 2.5.2 Sensitivity analysis

To systematically investigate the influence that each of the ten parameters assumed to be known has on the results of the base model (i.e. without interventions), we conduct a sensitivity analysis. For this, we fix all parameters except one at their assumed mean value (Table 1) and vary the free parameter across a range from 90% to 110% of its mean value to see the magnitude of effect that specific parameter has on the four outcomes of interest, see Figures in Section E of the Appendix. Table 3 shows the percentage change in each outcome of interest if we replace the mean of the free parameter with *±* 10% of the investigated parameters. Note, that the variation of 10% was chosen arbitrarily and should not be confused with the associated uncertainty of each parameter. A sensitivity analysis for the parameter *γ*, the rate of moving from the acute to the chronic compartment, is not shown because that parameter is fixed by our definition of the acute phase.

**Table 3:** Summary of sensitivity analysis showing percentage change in time until diagnosis, *R*^in^, total number of undiagnosed and true incidence as each quantity is changed from 90% to 110% of the mean value.

| Varied parameter |  | Time until diagnosis | Effect on: |  |  |
| --- | --- | --- | --- | --- | --- |
| | | | $R^{\text{in}}$ | # undiagnosed | Incidence |
| $N$ | Active PWID population size | 91% | 0% | 153% | 22% |
| $\mu$ | Mortality rate for PWIDs | 0% | 0% | 0% | 0% |
| $\mu + \rho$ | Mortality rate for HCV-infected PWIDs | 2% | 0% | -3% | -5% |
| $c$ | Permanent injection drug use cessation rate | -34% | 0% | -47% | -14% |
| $p$ | Clearance probability | 0% | 0% | 0% | 0% |
| prev | Prevalence among active PWIDs | 91% | 161% | 153% | 22% |
| $Y_A$ | Diagnosed acute cases | -6% | 0% | -5% | 0% |
| $Y_C$ | Diagnosed chronic cases | -42% | 1% | -47% | 1% |
| $Y_I$ | Diagnosed imported chronic cases | 0% | -1% | 0% | -1% |

#### 2.5.3 Scenario analyses

To illustrate how the model can be used to provide qualitative insights regarding the effect of different strategies on the disease dynamics, we created a set of eight scenarios (Table 4).

**Table 4:** Scenario analyses for interventions depending on treatment rates for active PWIDs not enrolled in NEPs (*τ_a_*), former PWIDs (*τ_f_*) and PWIDs in NEPs (*τ_e_*), enrollment rate into NEP (*ε*), relative susceptibility/infectiousness in NEP (*α*), permanent injection drug use cessation rate in a NEP (*c_e_*) and diagnosis rate (*r_e_*). The resulting prevalence among active PWIDs, the true incidence, the number of undiagnosed active and former PWIDs and the total number infected (i.e. undiagnosed and diagnosed, active and former) relative to the first scenario are reported. All rates are per year.

| Scenarios | Treatment |  |  | Needle exchange program (NEP) |  |  |  | Results |  |  |  |  |  |
| --- | --- | --- | --- | --- | --- | --- | --- | --- | --- | --- | --- | --- | --- |
| | # | $\tau_a$ | $\tau_f$ | $\tau_e$ | $\varepsilon$ | $\alpha$ | $c_e$ | $r_e$ | Prevalence among active PWIDs | True incidence | #undiagnosed active PWIDs | #undiagnosed former PWIDs | total # infected |
| No treatment, no NEP | 1 | - | - | - | - | - | - | - | 0.82 | 2146 | 10905 | 8389 | 83035 |
| Treatment, no NEP | 2 | - | 1 | - | - | - | - | - | 0% | 0% | 0% | 0% | -63% |
|  | 3 | 1 | - | - | - | - | - | - | -19% | 49% | 47% | 36% | -17% |
| No treatment, NEP | 4 | - | - | - | 0.1 | 0.5 | 0.056<sup>a</sup> | 0.072<sup>b</sup> | -18% | -19% | -18% | -14% | -16% |
|  | 5 | - | - | - | 0.1 | 1<sup>c</sup> | 0.2 | 0.072<sup>b</sup> | -15% | -10% | -31% | 22% | -8% |
| Treatment with increased diagnosis rates | 6 | - | - | 1 | 0.1 | 1<sup>c</sup> | 0.056<sup>a</sup> | 4 | -64% | 40% | -64% | -49% | -57% |
|  | 7 | - | - | 1 | 0.2 | 1<sup>c</sup> | 0.056<sup>a</sup> | 4 | -89% | -45% | -90% | -69% | -79% |
| Treatment, NEP | 8 | - | 1 | 1 | 0.1 | 0.5 | 0.056<sup>a</sup> | 4 | -82% | -49% | -80% | -61% | -91% |

<sup>a</sup> Value corresponds to the estimate for the cessation rate $c$ in the base model.
<sup>b</sup> Value corresponds to the estimate for the diagnosis rate $r_C$ in the base model.
<sup>c</sup> Value corresponds to no reduction in infectiousness and susceptibility compared to the base model.

The baseline scenario (Scenario 1) reflects the situation in which neither treatment nor NEPs are in place.

Scenarios 2 and 3 represent situations in which diagnosed PWIDs (former and active, respectively) are treated but no NEPs are in place.

In Scenarios 4-5 we explore the potential benefits of NEPs when no DAA treatment is available by independently varying each of the NEP-specific parameters. In Scenario 4 we assume a decrease in infectiousness and susceptibility (due to reduced needle sharing) from *α* = 1.0 to *α* = 0.5. In Scenario 5 we assume that NEPs contribute to early injection drug use cessation by reducing the average time spent injecting drugs from 17.8 years (*c_e_* = 0.056 [yrs^*−*1^]) to 5 years (*c_e_* = 0.2 [yrs^*−*1^]).

With Scenarios 6 and 7 we illustrate how the model can be used to study the effect of increased HCV testing by healthcare facilities where PWID can receive treatment but do not receive the benefits of NEPs. In these cases, the healthcare facilities are represented by *α* = 1 (no reduction in relative susceptibility/infectiousness) and *c_e_* = *c* = 0.056 [yrs^*−*1^] (no change in injection drug use cessation rates) as in the base model. In Scenario 6 we compare the effect of increasing the diagnosis rate from *r_C_* = 0.072 [yrs^*−*1^] in the base model (corresponding to an average of 14 years after infection) to *r_e_* = 4 [yrs^*−*1^], which corresponds to an HCV-infected individual being diagnosed, on average, 3 months after joining a healthcare facility due to regular health check-ups (this time period was based on personal communication with Martin Kåberg at the Stockholm NEP). We chose the enrollment rate into the NEP or healthcare facility to be *ε* = 0.1 [yrs^*−*1^] meaning that active PWIDs join, on average, 10 years after injection drug use initiation. In Scenario 7 we investigate the effect of increased diagnosis rates combined with greater availability of healthcare facilities by decreasing the average time until enrollment in healthcare from 10 years to 5 years after after injection drug use initiation, i.e. *ε* = 0.2 [yrs^*−*1^].

In Scenario 8 we demonstrate a combination of treating both active PWIDs in NEPs as well as former PWIDs; infectiousness is reduced and diagnosis rates are increased. For all treatment scenarios described above we chose a treatment rate of 1 [yrs^*−*1^], meaning individuals are successfully treated one year after HCV diagnosis, on average.

For our scenario analyses, we initialize the ODE system at the intervention-free endemic level for each parameter vector as calculated in Section 3.1 and then simulate the trajectory forward in time until the system has equilibrated in the steady state with interventions, see Figure 5 in the Appendix for an example. For this we use the trajectory function implemented in the R-package pomp (King et al., 2016). From this new equilibrium we compute the resulting HCV prevalence among active PWIDs, true annual incidence, number of undiagnosed among active and former PWIDs and total number of infected (i.e. undiagnosed and diagnosed, active and former); we report the corresponding deviation from the mean of the baseline situation, see Table 4. For simplicity we ignore drop-out from the needle exchange programs in all scenarios above but all of the intervention-specific parameters can be further explored and varied in the Shiny app.

**Figure 5:**
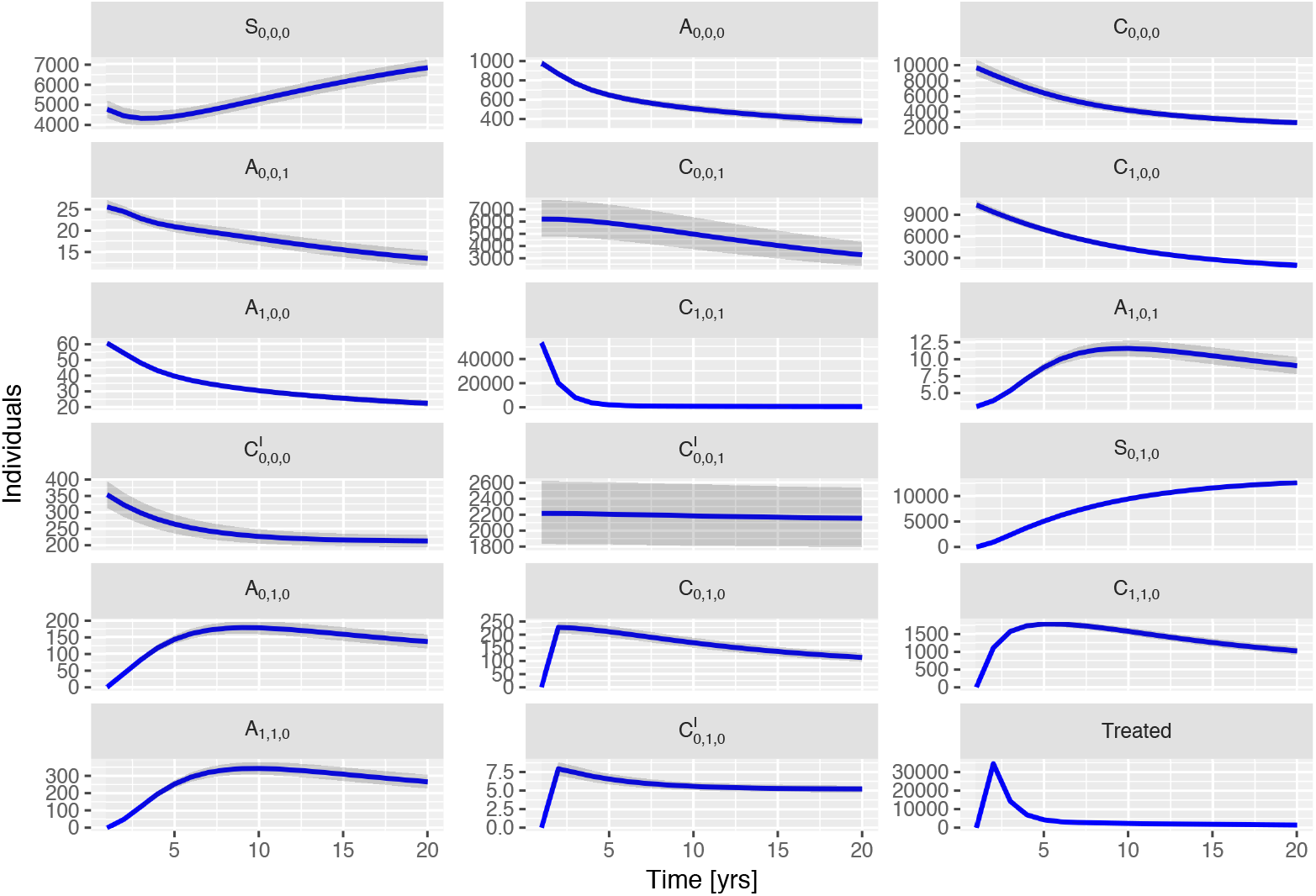
Solution of the ODE system including the number of treated for Scenario 8 initialized at the intervention free endemic level for 20 years into the future. In blue is the mean of all simulations and in grey shading the 95 % confidence interval.

#### 2.5.4 User interface: Shiny app

To complement our model and enable its use in public health practice, we developed an interactive platform through which the user can obtain model estimates for individual choices of parameters and instantaneously explore the sensitivity of the results to parameter changes. All outcomes presented throughout the manuscript can be obtained for user-specific sets of values by modifying all parameters assumed fixed along with their respective associated uncertainties. Additionally, the app provides visualization of each obtained parameter distribution and the sample size can be varied. The tool is made available in Stocks (2018) and a screenshot is shown in Figure 4 in the Appendix.

## 3 Numerical illustration

To demonstrate our model, we give a numerical illustration and estimate the number of undiagnosed cases, the true incidence, the time until diagnosis and the reproduction numbers in an intervention-free situation assuming endemic equilibrium. We base our calculations on surveillance data, i.e. the number of annual diagnosed cases, from Sweden, where HCV is endemic and the availability of NEPs has been, until recently, rather limited. When available, we use parameter estimates for Sweden and use other literature-informed values as needed (Table 1). Therefore, this section is meant as a proof of concept to illustrate our methods and not as a basis for public health decision-making; however, once all parameter estimates become available, our tool provides the infrastructure to monitor the key outcomes and intervention strategies as described above.

There is no uncertainty associated with the parameter *γ* because the length of the acute period is defined as six months (Mondelli et al., 2005). Moreover, in absence of a better alternative, we assume that the factor by which mortality of PWIDs increases due to chronic HCV is zero, however, the Shiny app allows for an increased mortality rate.

### 3.1 Key outcomes for intervention-free model in endemic state

Based on a sample of size 1000 and using the parameter estimates (Table 1), the annual true HCV incidence in an intervention-free situation in the endemic state is estimated to be 2146 (sd=35) cases, with an estimated average of 9.2 (sd=0.5) years between infection to diagnosis (see Figure 2). The estimated reproduction number *R*^in^ is 5.3 (sd=0.2) and *R*^out^ is 7.1 (sd=0.3) (see Figure 2).

The distribution of the number of undiagnosed cases in endemic state stratified by injection drug use status and stage of infection can be found in Table 2.

### 3.2 Sensitivity analysis: Baseline situation without interventions

Table 3 shows that time until diagnosis, the number of undiagnosed individuals and incidence are positively correlated with PWID population size and HCV prevalence and they are negatively correlated with time until permanent injection drug use cessation and number of chronic cases. The reproduction number *R*^in^ depends almost exclusively on prevalence: the higher the prevalence, the higher the reproduction number. We conclude that, based on the parameter values we investigated, the parameters for population size of active PWIDs, cessation rate, prevalence and number of diagnosed chronic cases are the most influential on the estimates; therefore, an increase in certainty for those four parameters would most effectively increase the precision of our results.

### 3.3 Impact of interventions

We investigated eight potential intervention scenarios and compared the results of four measures (prevalence among active PWIDs, incidence, number of undiagnosed cases and total number infected) to the baseline scenario (Scenario 1), see Table 4.

By model design, in a scenario in which treatment is available but no NEPs are in place, treating only former PWIDs (Scenario 2) results in a substantial reduction in the total number of infected individuals compared to the baseline but has no effect on prevalence, incidence or the number of undiagnosed individuals. In contrast, treating only active PWIDs (Scenario 3) additionally reduces prevalence but results in an increase in both incidence and the number of undiagnosed compared to the baseline scenario. This counter-intuitive effect arises because incidence is proportional to the product of prevalence and number of susceptible (Equation (5)): it increases when the number of susceptible individuals increases while prevalence is not reduced enough to compensate for this effect. Once re-infected, the PWIDs move from the susceptible to the undiagosed compartments where no treatment is available and remain there until diagnosis. As the diagnosis rate is low in the base model, few individuals proceed into the diagnosed compartments and, therefore, the number of undiagnosed increases.

In a situation where no treatment of PWIDs is available but NEPs are in place, reducing infectiousness and susceptibility to the disease, results in a reduction in all outcomes (Scenario 4). If injection drug use cessation rates increase, e.g. due to counselling in NEPs, this results in a reduction in all outcomes except for the number of infected former PWIDs (Scenario 5).

In Scenarios 6-7, treatment is made available only to PWIDs enrolled in healthcare facilities (hence, no reduction of infectiousness and susceptibility due to needle exchange). The effect of increased diagnosis rates in those centers by, e.g., regular health check-ups (Scenario 6), results in a reduction in prevalence, number of undiagnosed individuals and total number of infected, but an increase in incidence due to reinfection of treated individuals. In our example, incidence can be reduced by, e.g., increasing the enrollment rate into the healthcare facilities (Scenario 7).

Combining treatment of former PWIDs and treatment in NEPs (Scenario 8) leads to a reduction in all outcomes.

## 4 Discussion

We developed and implemented a model to help enable countries to monitor progress towards the WHO goal of viral hepatitis elimination, and to evaluate how needle exchange programs and DAA treatment may influence this progress. Within the scope of the WHO elimination targets, this is highly relevant for public health. We demonstrated the model and the output it provides using an illustrative data set. In addition, we converted the model into an accessible, web-based tool intended for public health professionals.

For the first objective (monitoring), the model uses several country-specific parameters (HCV prevalence, active PWID population size, reported HCV incidence, PWID-specific death-rate, and permanent cessation rate) as well as certain HCV-specific parameters (duration of acute period and spontaneous clearance probability). Based on these input parameters, the model is able to generate four key outcomes the WHO recommends be monitored that describe the current HCV situation in the target country, namely the number of undiagnosed PWIDs, the true incidence, the average time until diagnosis and the reproduction numbers.

PWID-specific parameters are often accompanied by high uncertainty; therefore, our model allows this uncertainty to be quantified. The uncertainty in the input parameters is then translated into uncertainty in the output parameters by Monte Carlo methods, which results in the key outcomes being distributions rather than point masses. Furthermore, our modelling framework provides sensitivity analyses based on the specified input parameters in order to investigate which parameter values are most influential on the estimated outputs. This can help countries to prioritize which parameters should be the focus of further public health investigations to most effectively increase precision of the results (e.g., Table 3). For example, the dataset we investigated, the analysis showed that increased certainty regarding three key quantities (prevalence among active PWIDs, PWID population size, and average duration of injection drug use) would most improve the precision of the results and should be investigated further.

For the second objective (intervention analysis), the effect of different, country-specific interventions on prevalence, incidence, number of undiagnosed and total number of infected can be investigated. Here, the model offers great flexibility in terms of which scenarios can be explored: as illustrated in Table 3, treatment rates depending on injecting drug use status and access to health care facilities and NEPs can be varied, the effect of increased diagnosis rates due to regular healthcare checkups in NEPs or health care centers can be investigated and availability of and drop-out from those facilities can be quantified. Additionally, the model allows for the explicit investigation of two NEP-specific benefitis that can potentially reduce HCV transmission in the PWID population. First, through one of the main functions served, NEPs can reduce HCV transmission by providing access to clean needles and injection equipment. Second, if connected with counseling and/or opioid substitution therapy programs, NEPs have the potential to increase injection drug use cessation rates. A strength of our model is that these effects can be investigated in isolation or in combination, which can reveal how these different factors interplay.

In addition, our scenario analyses showed the importance of stratifying individuals by their injection drug use status. For example, our model produced rather counter-intuitive findings, with respect to incidence and the number of undiagnosed individuals, when we considered a potential scenario in which treatment was provided to active PWIDs in the absence of NEPs and with relatively low rates of diagnosis. However, our model was able to show greater changes in outcomes relative to baseline for scenarios that combined treatment with increased diagnosis rates compared to those that did not change diagnosis rates; this was only possible because we stratified by diagnostic status. This stresses the importance of considering diagnosis rates explicitly in the modelling process and is further supported by studies reporting the importance of HCV screening, see e.g. Scott et al. (2017a, 2018a,b). Models that do not differentiate cases by diagnostic status may fail to accurately estimate the success of treatment interventions because they ignore that undiagnosed infected individuals do not benefit from treatment while, at the same time, these individuals contribute to the infectious pressure in the population. The ability to uncover these relationships is a strength of our model and illustrates the role models can play in potentially preventing costly but inefficient, or even counter-effective, interventions. Making our model available as an open software tool gives countries the ability to test various potential intervention strategies beforehand, in an effort to prevent potentially negative real-world consequences and use limited resources most effectively.

There are, of course, certain limitations to our modelling approach and complexities not considered. First, we assumed homogeneous mixing of needle sharing among PWIDs; however, in reality, needle sharing behavior can be heterogeneous (Strathdee et al., 2001). Therefore, the model could be made more complex by considering a spatial and social structure and inclusion of distinct risk groups that differ with respect to needle sharing rates. Second, to simplify our model, we assumed injection drug use cessation to be permanent, however, relapse is common (Galai et al., 2003). As more data become available, the model could be easily extended to include estimated relapse rates. Third, we limited our interventions to treatment and NEPs and did not consider opiate substitute therapy programs. We chose not to incorporate these programs explicitly in our model because they mainly affect the transition rate between active and former PWIDs, which can be indirectly modelled by increasing cessation rates, which our model already allows. We also assumed that initial infection and re-infection occur at the same rate; however, a change in injection drug use behavior following HCV infection and treatment is possible, but was not considered here. Finally, a general limitation of compartmental models is that they implicitly assume that the time in each compartment is exponentially distributed, which might not always be realistic – distributions with more pronounced modes could be achieved by dividing the compartments into several sub-classes.

Recently and independent of our work, models have been proposed presenting similar stratifications as we have used here (Scott et al., 2017a,b, 2018a,b). Some are much more detailed with respect to stratification by age, cascade of care and disease progression; however, none account for NEPs explicitly. The analyses of Scott et al. often focus on the impact of testing in a specific country/prevalence setting; on the other hand our model is less detailed in some of these aspects but offers more flexibility as it is meant as an open software tool for any high income country where the main route of transmission is injection drug use among active PWIDs. As such, availability of country-specific parameter estimates is important. By keeping our model relative simple, e.g. with respect to disease progression and age structure, we limit its dependence on difficult-to-obtain parameters and, at the same time, strive to ensure that model inputs and outputs are easily interpretable.

We expect that our model can be used as a tool for intervention and treatment planning, allocation of resources (such as funding, time and personnel) and support for evidence-based public health decision-making aimed at reaching the WHO HCV elimination targets. To our knowledge, our model is the first to be developed into a ready-to-use tool that enables measurement of country-specific WHO indicators to allow planning and monitoring of progress toward the elimination of viral hepatitis.

## Acknowledgments

We thank Maria Axelsson, Viktor Dahl, Stephan Stenmark, Ann-Sofi Duberg and Martin Kåberg for their expertise and insights about HCV transmission in Sweden and Michael Höhle and Ka Yin Leung for helpful comments to our manuscript. TS and TB were supported by the Swedish research council (2015_05182_VR for TS and 2015_05015_VR for TB).

## A Transmission equations

The differential equations describing our model are given as follows. Note that this is a deterministic system so the state variables are continuous rather than integer-valued.

### 1. PWIDs in the base model

a) Active PWIDs:

(i) Undiagnosed

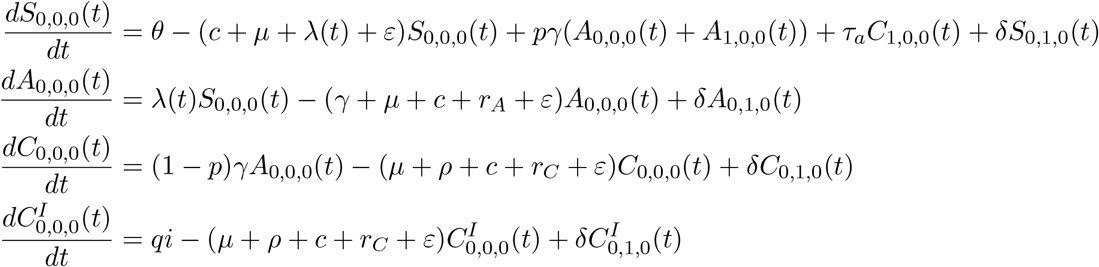
(ii) Diagnosed

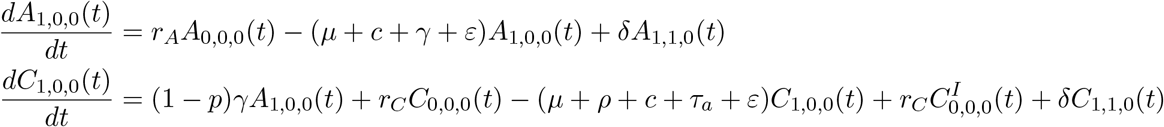
b) Former PWIDs:

(i) Undiagnosed

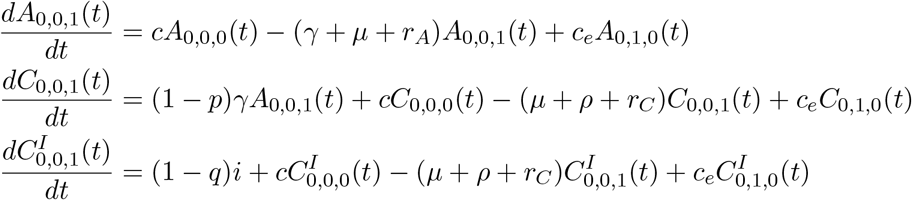
(ii) Diagnosed

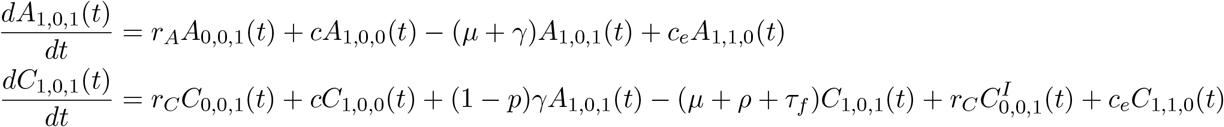

### 2. PWIDs enrolled in NEPs

(i) Undiagnosed

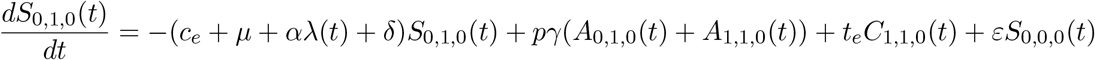

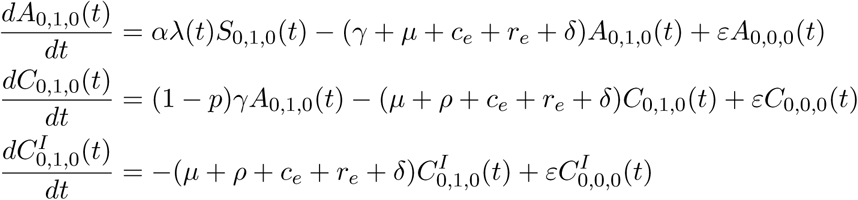
(ii) Diagnosed

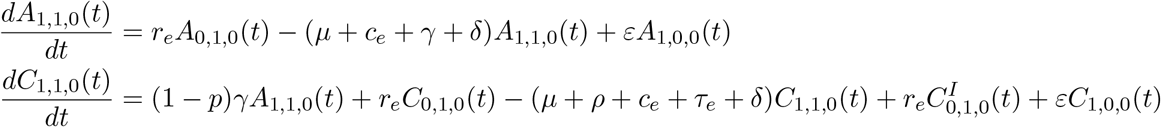

All initial values are fixed at the endemic level without interventions as described in Section 2.5. Moreover, the initial values satisfy

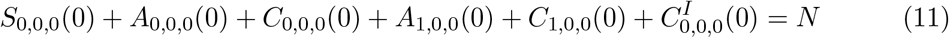

for the compartments in the base tranmission model and

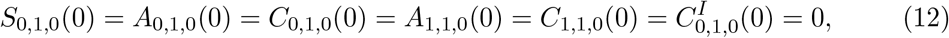

for the compartments in the NEP since we assume that initially no NEP is implemented. If the enrollment rate is *ε >* 0 the values in Equation (12) change over time. Since the population size of active PWIDs, *N*, is assumed to be constant over time, the influx rate into the susceptible compartment is given as

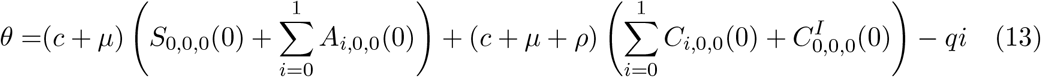

which is the accumulated rate of all individuals leaving the active PWIDs community by death or injection drug use cessation each year minus the rate of individuals entering the community by importation, *q · i*, every year. Moreover, we chose the fraction of active imported chronic cases, *q*, equals the fraction of time an individual spends, on average, in the active, chronically-infected compartment 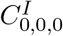. If *t_ac_* denotes the average time an active PWIDs is chronically infected and *t_fc_* the average time a former PWIDs is chronically-infected then this translates to

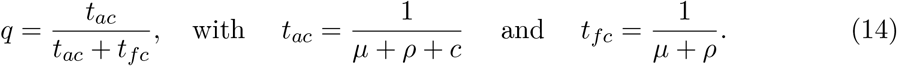

## B Conditional times until diagnosis

The average time until diagnosis given the individual is diagnosed in compartment 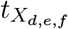 with *X* ∈ {*S, A, C*} can be calucalted as

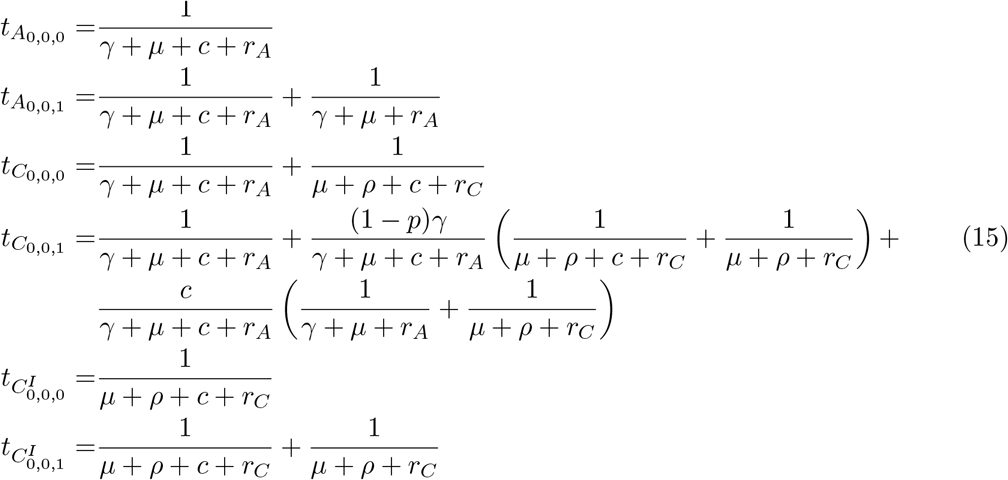

## C Reproduction number *R^in^* without imported cases

## D Screenshot shiny app

## E Visualization of the sensitivity analysis

The following plots visualize the results of the sensitivity analysis in Table 3. They show the resulting change in the four key outcomes if one input parameter is varied from 90% to 110% of its mean, respectively, with all other input parameters assumed fixed.

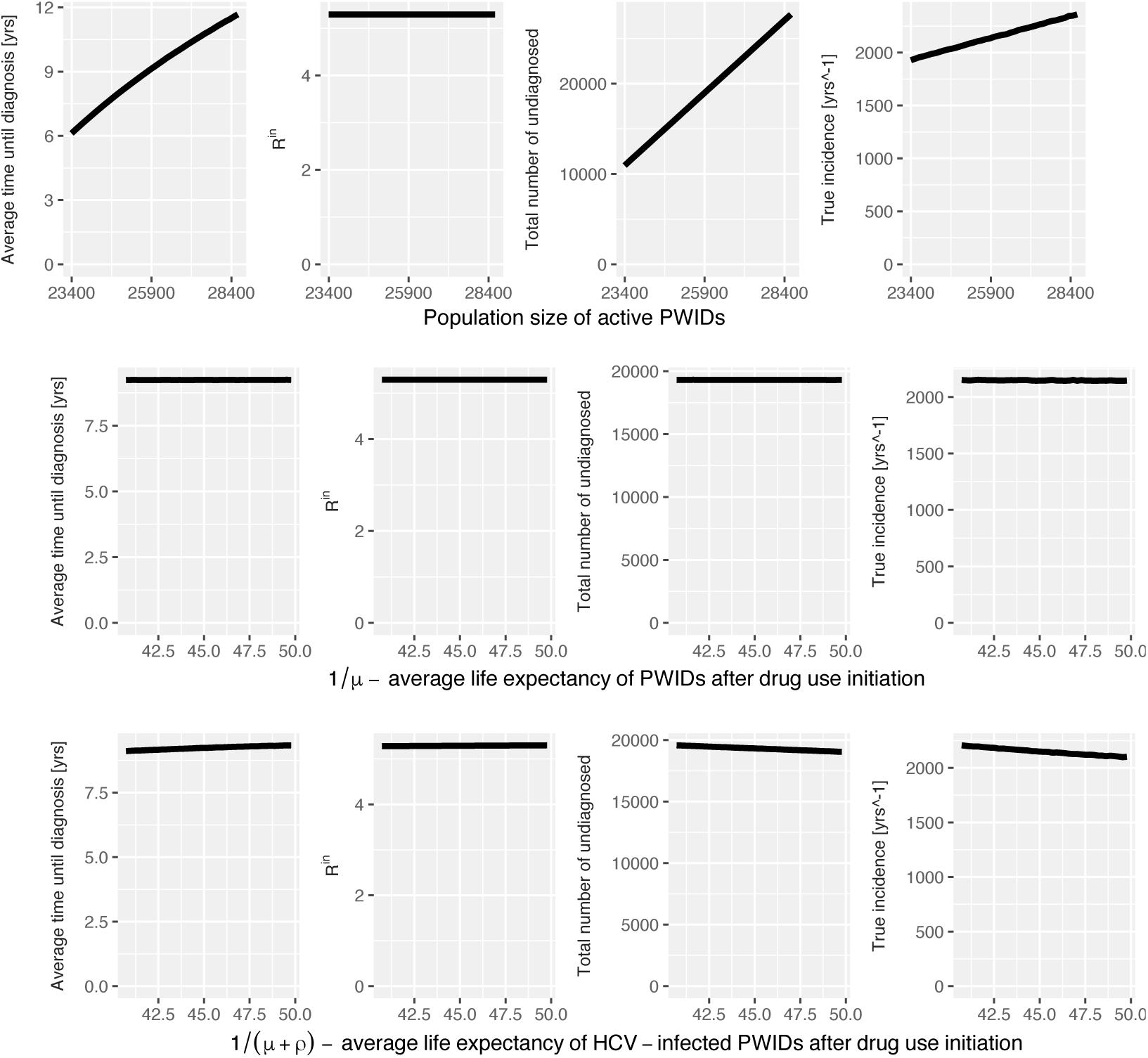

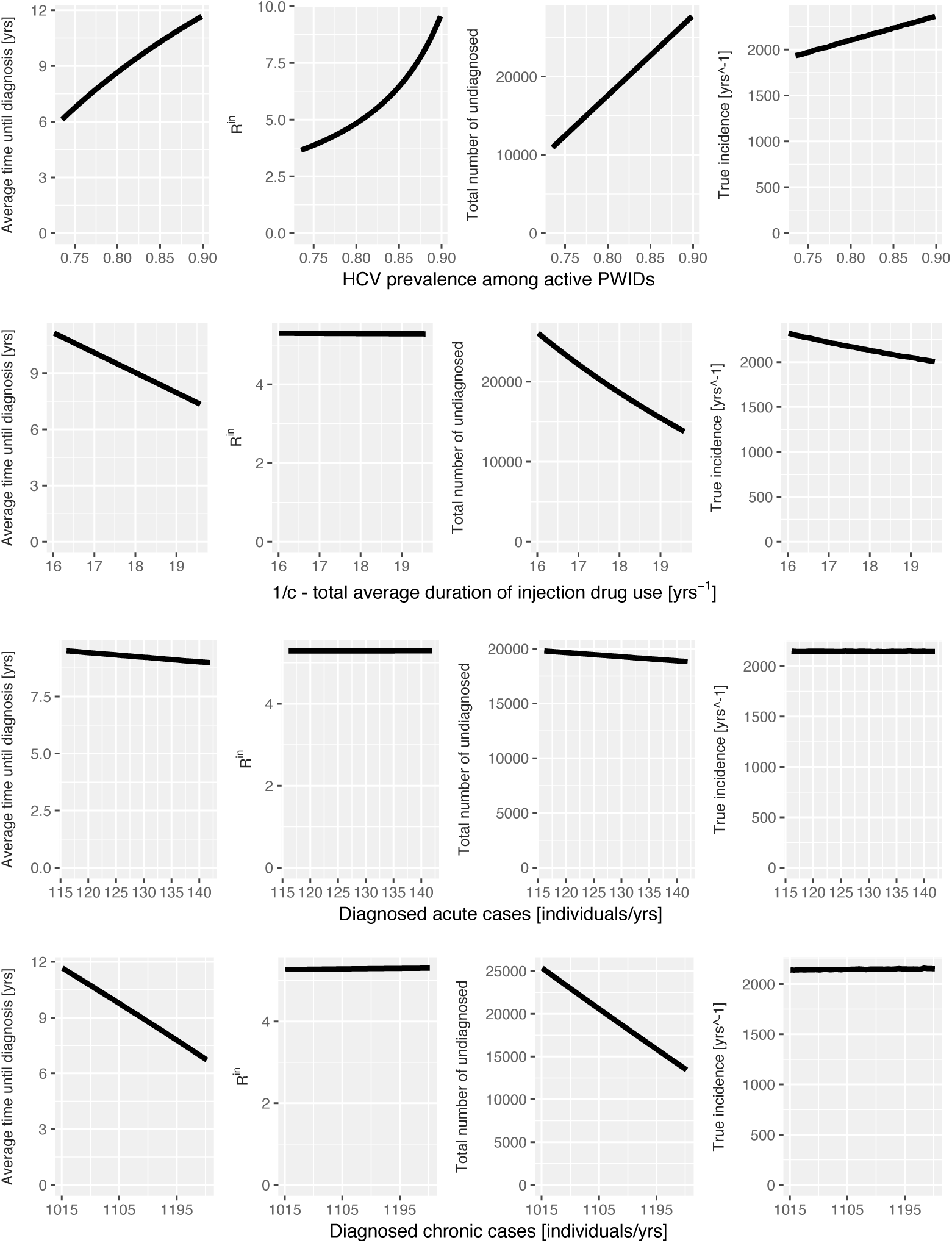

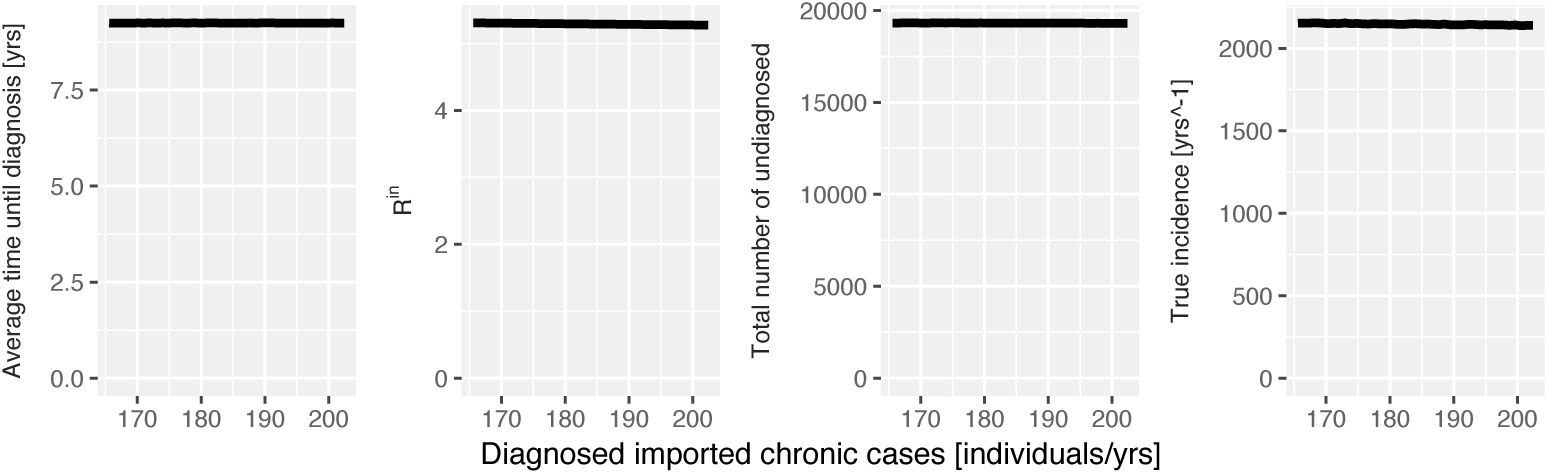

## F Simulation of trajectories

